# Glycan remodeled erythrocytes facilitate antigenic characterization of recent A/H3N2 influenza viruses

**DOI:** 10.1101/2020.12.18.423398

**Authors:** Frederik Broszeit, Rosanne J. van Beek, Luca Unione, Theo M. Bestebroer, Digantkumar Chapla, Jeong-Yeh Yang, Kelley W. Moremen, Sander Herfst, Ron A.M. Fouchier, Robert P. de Vries, Geert-Jan Boons

## Abstract

During circulation in humans and natural selection to escape antibody recognition for decades, A/H3N2 influenza viruses emerged with altered receptor specificities. These viruses lost the ability to agglutinate erythrocytes critical for antigenic characterization and give low yields and acquire adaptive mutations when cultured in eggs and cells, contributing to recent vaccine challenges. We examined receptor specificities of A/H3N2 viruses, revealing that recent viruses compensated for decreased binding of the prototypic human receptor by recognizing 2,6-sialosides on extended LacNAc moieties. Erythrocyte glycomics showed an absence of extended glycans, providing a rationale for lack of agglutination by recent A/H3N2 viruses. A glycan remodeling approach installed functional receptors on erythrocytes, allowing antigenic characterization of recent A/H3N2 viruses and confirming the cocirculation of several antigenically different viruses in humans. Computational studies of HAs in complex with a sialoside having an extended LacNAc moiety revealed that mutations distal to the RBD reoriented the Y159 side chain, resulting in an extended receptor binding site.

## Main

Human influenza A viruses have a remarkable ability to evolve and evade neutralization by antibodies elicited by prior infections or vaccinations ^3^. This antigenic evolution, or drift, is mainly caused by amino acid substitutions in the globular head of the hemagglutinin (HA) protein where binding occurs with sialic acid receptors of host cells ^4,5^. These substitutions in circulating influenza viruses lead to antigenic differences as compared to employed vaccines, resulting in poor vaccine-mediated protection ^6^. Therefore, the World Health Organization (WHO) Global Influenza Surveillance and Response System (GISRS) continuously monitors antigenic changes in circulating influenza viruses and recommends updated compositions of influenza vaccines biannually ^7^.

Antigenic surveillance and vaccine strain selection rely predominantly on the hemagglutination inhibition (HI) assay, in which the ability of serum antibodies to block receptor binding by the influenza virus HA protein is quantified ^8^. Such antibodies prevent virus-mediated agglutination of erythrocytes and are measured as a correlate of protection. The HI assay makes it possible to select virus strains that are antigenically representative of circulating viruses for vaccine development.

A/H3N2 viruses, which are associated with high morbidity and mortality, exhibit a particularly rapid antigenic drift, and as a result the WHO has recommended 28 vaccine strain updates since these viruses started circulating in humans in 1968 ^9^. In recent years, the rapid antigenic evolution of A/H3N2 viruses coincided with altered receptor usage, which in turn has resulted in their inability to agglutinate fowl erythrocytes. As a result, antigenic characterization of circulating A/H3N2 viruses using the HI assay is increasingly difficult, complicating the selection of appropriate vaccine strains. The receptor-binding phenotype of recent A/H3N2 viruses is also hampering virus replication under laboratory conditions for amplification of clinical isolates, and leads to adaptive substitutions when grown in embryonated chicken eggs ^10^. The difficulties to antigenically characterize circulating A/H3N2 viruses, in particular those belonging to the dominant 3C.2a clade, and the inability of large-scale virus production without egg-adaptation has led to serious problems with A/H3N2 influenza vaccine production and effectiveness.

HAs of human influenza viruses bind to cell surface glycans carrying terminal *α*2,6-linked sialic acid moieties (Neu5Ac(α2,6)Gal) ^11^. These receptors are usually part of *N*-linked glycans, which are highly complex biomolecules composed of a core pentasaccharide modified by various numbers and patterns of branching *N*-acetylglucosamine (GlcNAc) moieties ^12^. These branching points can be extended by several *N*-acetyl-lactosamine (Gal(β1,4)GlcNAc, LacNAc) repeating units, which in turn can be capped by various types of fucosylation and sialylation. Several studies have shown that not only the *α*2,6-linked sialoside but also the underlying oligosaccharide structure of *N*-linked glycans can contribute to HA binding selectivities ^13-17^. Antigenic pressure results mainly in amino acid substitutions near the receptor binding site of HA, which may result in changes in glycan receptor specificities. Thus, failure of contemporary A/H3N2 viruses to agglutinate erythrocytes, the poor replication in mammalian cells and the emergence of egg adaptive mutations are probably due to an adaptation to glycan receptors that are not expressed by these cell substrates. Understanding of the evolution of receptor usage by A/H3N2 viruses at a molecular level will open avenues to address challenges in surveillance and vaccine production for these viruses.

We have examined receptor specificities of A/H3N2 viruses representing different evolutionary time points and clades using a novel glycan microarray and identified the minimal receptor for recent, non-agglutinating A/H3N2 strains. Glycan analysis of cell surface *N*-glycans on various erythrocytes revealed an absence of such receptors. We developed an exo-enzymatic cell surface glycan remodeling strategy to install appropriate receptors on fowl erythrocytes to regain binding by contemporary A/H3N2 viruses. Such viruses agglutinated the glyco-engineered erythrocytes and made it possible to antigenically characterize A/H3N2 viruses by HI assays. It revealed substantial antigenic differences of circulating virus isolates to current vaccine strains providing a rationale for poor vaccine performance.

## Results

### Glycan microarray analysis to determine A/H3N2 receptor specificity

Although glycan microarray technology has been used to examine receptor requirements of HAs ^18^, these were not populated with biologically relevant glycans to establish minimal receptor requirements. This information is, however, critical to understand how receptor binding has evolved over time and how a lack of expression of specific glycans by erythrocytes or laboratory hosts may have resulted in a loss of agglutination or a lack of propagation, respectively. We have constructed a glycan array that is populated with biologically relevant bi-antennary *N*-glycans having different numbers of LacNAc repeating units in various structural configurations. These compounds are either unmodified (compounds **1**-**3**), capped by avian *α*2,3-linked (compounds **4**-**6**) or human *α*2,6-linked sialosides (compounds **7**-**17**). Most naturally occurring *N*-linked glycans have asymmetrical architectures in which the various antennae are modified by oligo-LacNAc moieties of different lengths ^19^. To probe the importance of glycan architecture for HA recognition, we employed a chemo-enzymatic strategy to prepare α2,6-sialosides having such architectures (**8, 9, 11, 12, 13, 15, 16** and **17**). These compounds made it possible to probe the importance of mono- *vs*. bidentate binding interactions, preference for a specific antenna, and possible interference of a neighboring arm. The structures on the array represent typical *N*-linked glycans found in human respiratory tissue making it relevant for HA binding studies ^20^. The quality of the printing was validated by probing the array with the lectins ECA, SNA and MAL1 (Fig. 1A and Fig. S1).

**Figure 1.**
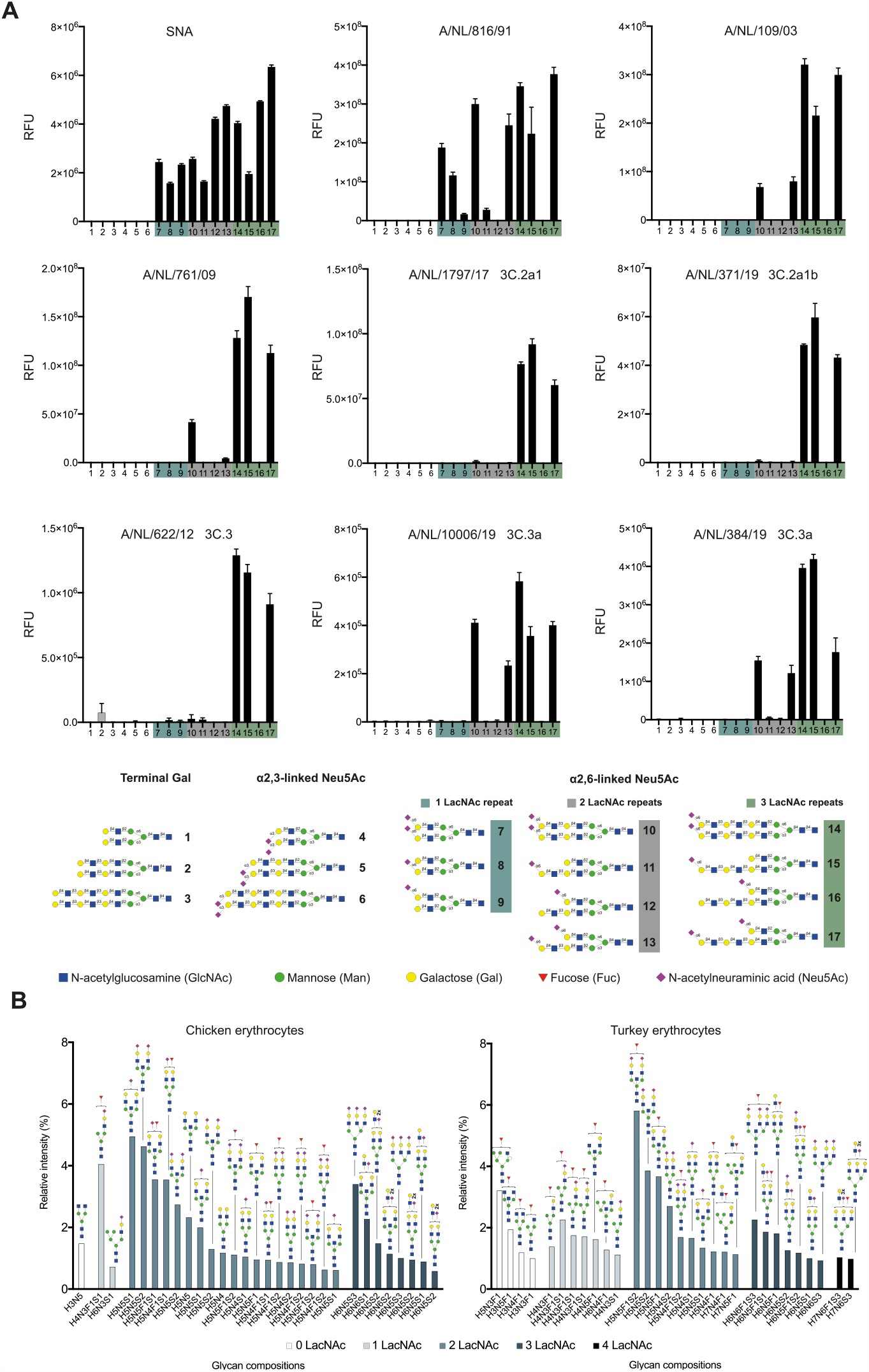
**(A) Receptor binding specificities of representative A/H3N2 viruses using glycan microarray analysis**. Binding was visualized using a human anti-H3 stalk antibody (CR8020). Bars represent the mean ± SD of relative fluorescence units (RFU) **(B) Glycomic analysis of N-glycosylation of unmodified fowl erythrocytes**. The 30 most abundant N-glycans on erythrocytes from chicken and turkey (based on relative intensity, excluding high-mannose type N-glycans, for all structures refer to Data S1) sorted by abundance and number of LacNAc units. Proposed structures are assigned to detected glycan compositions.

The glycan array was probed with various A/H3N2 viruses, representing distinct evolutionary time points and clades and having different abilities to agglutinate erythrocytes (Fig. S2). It includes A/NL/816/91 (NL91) which can agglutinate chicken, turkey and guinea pig erythrocytes, A/NL/109/03 (NL03) which only agglutinates turkey and guinea pig erythrocytes ^21^ and A/NL/761/09 (NL09) which only agglutinates 2,6-resialylated turkey and guinea pig erythrocytes ^1,22,23^. During the past decade, A/H3N2 viruses have evolved into distinct, cocirculating antigenic groups, referred to as clades (Fig. S2). We examined A/NL/1797/17 (NL17) and A/NL/371/19 (NL19) as recent examples of the 3C.2a clade that insufficiently hemagglutinate all commonly used erythrocytes for HI assays, and poorly infect MDCK cells^24^. The 3C.3a clade is represented by A/NL/10006/19 and A/NL/384/19, and their evolutionary predecessor A/NL/622/12 (3C.3). Although 3C.3 viruses cannot agglutinate any erythrocyte type, 3C.3a viruses are unique as they regained an ability to agglutinate turkey and guinea pig erythrocytes (Fig. S2).

Whole viruses were applied to the microarray and detection of binding was accomplished by a human anti-H3 stalk antibody (CR8020) (Fig. S1). NL91 recognized most of the human-type receptors, including compounds that have an *α*2,6-sialoside on a mono-LacNAc residue (glycans **7**-**9**, Fig. 1A and Fig. S3). Compound **8** exhibited a substantial greater responsiveness compared to **9** indicating that this virus has a preference for a sialoside at the *α*1,3-arm. Interestingly, the sialyltransferase, ST6Gal1, which is solely responsible for installing human-type receptors, preferentially modifies the *α*1,3-arm of *N*-linked glycans ^25^. Compounds **7** and **8** did bind similarly demonstrating that an additional sialic acid at the *α*1,6-arm does not substantially contribute to binding. Another unanticipated observation was that compounds **12** and **16** did not exhibit binding whereas **9** showed responsiveness highlighting that an extended and unmodified LacNAc moiety at the *α*1,3-arm can block recognition of the other arm. Collectively, the results show that the minimal receptor for NL91 is a biantennary *N*-glycan having two LacNAc moieties modified by a single sialoside (glycan **8**).

NL03 and NL09 recognized far fewer glycans and did not bind to structures having their *α*2,6-sialosides at a mono-LacNAc moiety (**7-9, 12** and **16**). This observation indicates that the minimal receptor for these viruses is a bis-sialylated *N*-glycan having at least one di-LacNAc moiety (glycan **13**). NL17 and NL19 (3C.2a) showed only strong responsiveness to **14, 15** and **17**. These glycans have in common that at least one of the arms is extended by three consecutive LacNAc units that is further modified by an *α*2,6-sialoside. Thus, a glycan having four LacNAc units arranged in an asymmetrical manner (**15**) represents the minimal receptor for these viruses. Mono-sialylated derivative **15** gave a similar responsiveness compared to the bis-sialosides **14** and **17** indicating that a bidentate binding event does not substantially contribute to recognition as previously suggested ^15^. Instead, it appears that reduction in binding of an *α*2,6-sialyl LacNAc moiety, which is the proposed prototypic human receptor, has been compensated by recognition of sialosides at an extended LacNAc chain. 3C.3a viruses (A/NL/10006/19 and A/NL/384/19) exhibited a similar binding profile as NL03 and 09 and bound to bis-sialosides **10** and **13** having the Neu5Ac residue at a di-LacNAc chain. Interestingly, their ancestor (A/NL/622/12, 3C.3) required the Neu5Ac residue to be presented on a tri-LacNAc structure similar to the requirement of 3C.2a viruses. Thus, recent 3C.3a viruses have regained an ability to recognize shorter structures.

### Glycomic analysis of chicken and turkey erythrocytes

Next, we examined structures of *N*-linked glycans expressed by chicken and turkey erythrocytes and compared the data with the receptor requirements of the various A/H3N2 viruses. Membrane fractions of the cells were treated with PNGase F to release the *N*-glycans which were isolated by solid phase extraction using C18 and Porous Graphitized Carbon (PGC) cartridges, and then analyzed by liquid chromatography mass spectrometry (LC-MS) ^26^. The 30 most abundant complex type *N*-glycan compositions for the two cell types are presented in Fig 1B. Strikingly, chicken erythrocytes did not substantially express *N*-glycans with at least four LacNAc units, which is the minimal epitope requirement for contemporary non-agglutinating A/H3N2 viruses. Turkey erythrocytes did express some glycans with this number of LacNAc units, but the majority was assigned as tri- and tetra-antennary glycans because of substitution with three or four sialic acids. The latter was supported by selectively releasing bi-antennary *N*-glycans using Endo F2, and in this case LC-MS analysis did not detect glycans having four LacNAc moieties (Fig. S4). Thus, turkey erythrocytes also do not substantially display sialylated epitopes having three consecutive LacNAc moieties. Chicken erythrocytes expressed substantial quantities of high mannose glycans (Fig. S5) whereas turkey cells displayed almost none of these structures. The greater abundance of complex type glycans on turkey erythrocytes offers a possible rationale for the ability of the NL03 and NL09 A/H3N2 viruses to agglutinate unmodified or α2,6-resialylated turkey erythrocytes, respectively.

### Glycoengineering of erythrocytes to install functional cell surface receptors

We embarked on a strategy to enzymatically remodel glycans of fowl erythrocytes to install receptors for A/H3N2 viruses of the 3C.2 clade to make them suitable for HI assays (Fig. 2A). Treatment of erythrocytes with a neuraminidase was expected to remove sialic acids and reveal terminal galactosides which are appropriate acceptors for installing additional LacNAc moieties. The latter residues can be introduced by the concerted action of the enzymes B4GalT1 and B3GnT2, which sequentially install *β*1,4-linked galactoside and *β*1,3-linked *N*-acetyl-glucosamines, respectively. The terminal galactosides of the resulting extended LacNAc moieties can then be modified by the sialyltransferase ST6Gal1 to install terminal α2,6-linked sialosides ^27^. The enzymatic remodeling was conveniently performed by incubating the erythrocytes with the neuraminidase from *Arthrobacter ureafaciens* for 6 h after which B4GalT1, B3GnT2, ^28^ UDP-Gal and UDP-GlcNAc were added followed by incubation overnight. Next, the cells were pelleted by centrifugation, washed to remove the enzymes and sugar nucleotides and then incubated with ST6Gal1 in the presence of CMP-Neu5Ac for 4 h. Glycomic analysis of the resulting cells, which were denoted as 2,6-Sia Poly-LN cells, confirmed that the antennae of the *N*-linked glycans had been extended by additional LacNAc moieties, and both cell types expressed sialylated structures having four LacNAc units (Fig. 2B). Analysis of glycans on turkey erythrocytes released by Endo F2 treatment confirmed the presence of bi-antennary glycans that have four LacNAc moieties and are potentially suitable receptors for contemporary A/H3N2 viruses (Fig. S4). Chicken and turkey erythrocytes express a mixture of *α*2,3- and *α*2,6-linked sialosides. To examine whether an increase in the abundance of *α*2,6-sialosides would improve agglutination, control cells were prepared by treatment with neuraminidase and resialylation with ST6Gal1 (denoted as 2,6-Sia cells). As negative control, we employed cells that have extended LacNAc moieties but lack sialic acids (denoted as Poly-LN cells). Phenotypic properties of the glyco-engineered erythrocytes were examined using the hemagglutination (HA) assay (Fig. 2C). As expected, NL91 agglutinated unmodified, 2,6-Sia and 2,6-Sia poly-LN erythrocytes, which was in agreement with the finding that these viruses can employ *N*-glycans that have simple and extended α2,6-sialylated structures. NL03 agglutinated unmodified turkey erythrocytes, but interestingly also α2,6-resialylated chicken erythrocytes. The latter may be due to an increase in the abundance of α2,6-linked sialosides having two consecutive LacNAc repeats, which are present on chicken erythrocytes. α2,6-Resialylation of turkey erythrocytes was sufficient to recover agglutination of NL09, and in this case the greater abundance of α2,6-sialylation on extended structures already present on these cells is probably responsible for the improved agglutination. Importantly, NL17 and NL19 (3C.2a) agglutinated only erythrocytes that were enzymatically remodeled to have extended sialylated LacNAc moieties (2,6-Sia Poly-LN cells). Similar results were obtained for A/NL/622/12 (3C.3), which is in agreement with receptor requirements similar to viruses of the 3C.2a clade. As anticipated, the 3C.3a viruses (A/NL/10006/19 and A/NL/384/19), which have reverted to recognize shorter structures, could also agglutinate *α*2,6-resialyated turkey erythrocytes.

**Figure 2.**
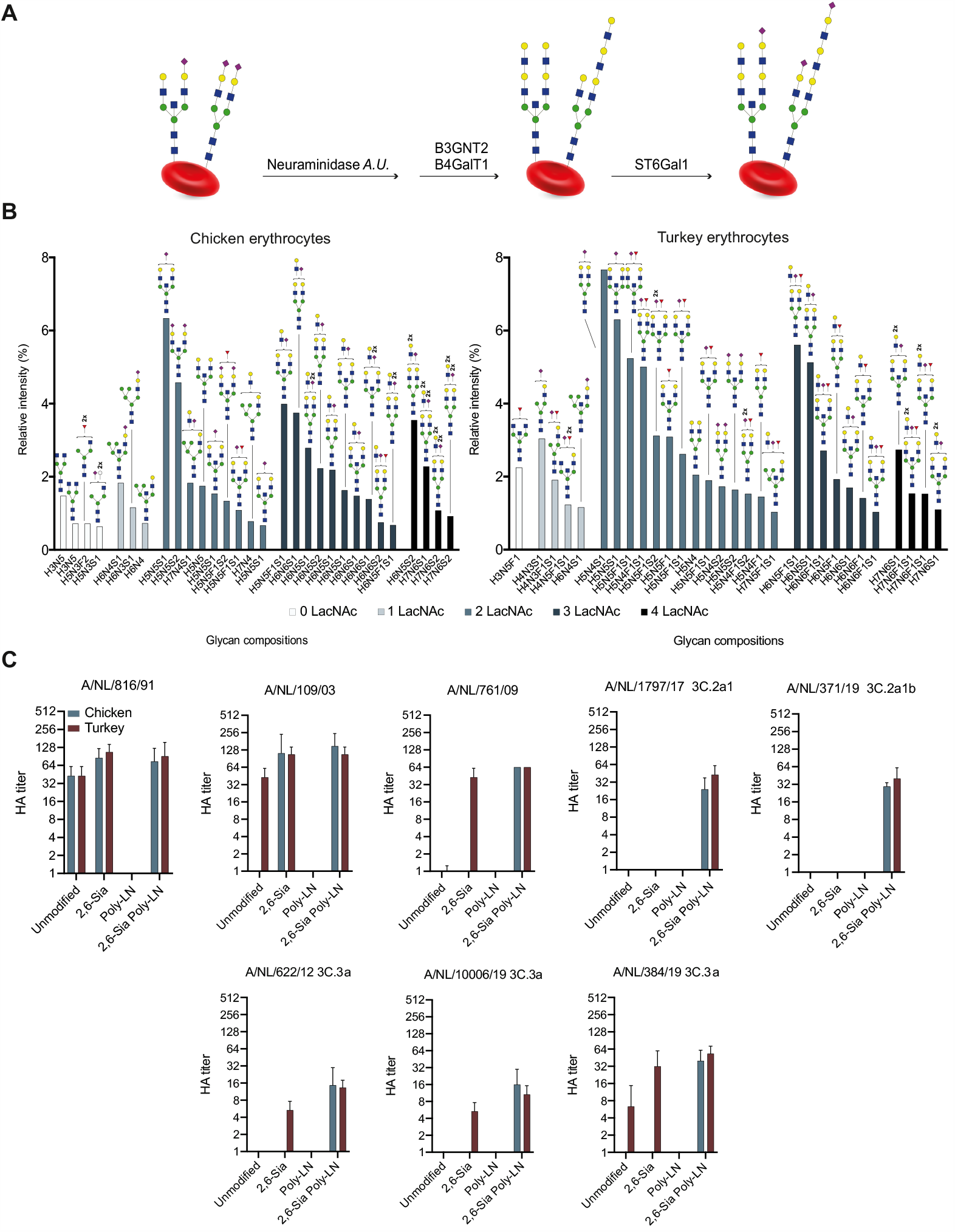
**(A) Schematic overview of enzymatic modification of erythrocytes**. Neuraminidase *A*.*U*.: Neuraminidase from *Arthrobacter ureafaciens*; B3GnT2: β-1,3-*N*-acetylglucosaminyltransferase 2; B4GalT1: β-1,4-galactosyltransferase 1; ST6Gal1: α-2,6-sialyltransferase 1. **(B) Glycomic analysis of N-glycosylation of enzymatically modified fowl erythrocytes**. The 30 most abundant N-glycans on the enzymatically modified erythrocytes from chicken and turkey (based on relative intensity, excluding high-mannose type N-glycans, for all structures refer to Data S2) sorted by abundance and number of LacNAc units. Proposed structures are assigned to detected glycan compositions. **(C) Hemagglutination assays with representative A/H3N2 viruses and modified erythrocytes**. A/NL/816/91, A/NL/109/03, A/NL/761/09, A/NL/1797/17, A/NL/371/19, A/NL/622/12, A/NL/10006/19 and A/NL/384/19 tested with modified erythrocytes (2,6-Sia Poly-LN) from chicken (blue) and turkey (red). Unmodified, 2,6 resialylated (2,6-Sia) and extended desialylated (Poly-LN) erythrocytes were added as controls. Assays were performed in full biological triplicates in the presence of oseltamivir and the means ± SEM were plotted.

Next, HA assays were performed with a collection of A/H3N2 viruses (Table S1) to validate the robustness of the glyco-engineering method, and the focus was on contemporary A/H3N2 viruses that have lost the ability to hemagglutinate unmodified erythrocytes and do not replicate efficiently in wild-type MDCK cells. Several recent vaccine strains heavily adapted to growing in eggs, an A/H1N1 and an influenza B strain were included as controls. As expected, pre-2000 A/H3N2 strains efficiently agglutinated unmodified chicken and turkey erythrocytes. Importantly, A/H3N2 viruses that emerged after 2010 only agglutinated the 2,6-Sia Poly-LN cells having extended sialylated epitopes. Although some contemporary A/H3N2 viruses can agglutinate turkey erythrocytes when applied undiluted, especially the two 2019 3C3a viruses, extended sialylated LacNAc moieties increased the efficiency of agglutination by 3 to 10-fold, including the A/H1N1 and influenza B controls. We performed a time course to determine the stability of the glyco-engineered cells, and no loss in titer or autoagglutination for up to three weeks was observed similar to unmodified cells (Fig. S6).

The 2,6-Sia Poly-LN cells were employed to antigenically characterize typical recent seasonal A/H3N2 viruses of various clades by HI assay using post-infection ferret sera (Table 1). All antisera showed robust inhibition of the homologous viruses and variable inhibition of heterologous viruses. Egg-derived vaccine strains displayed poor correspondence with data generated with cell-passaged viruses of the same clade. Antisera raised against cell-passaged virus isolates generally showed greater clade specificity compared to egg-derived vaccine strains and vaccine strains showed generally broader sensitivity to inhibition than observed with cell-passaged viruses. As a consequence, the HI assay revealed that antisera raised against recent vaccine viruses, including A/Kansas/14/17 that was selected for the 2019/2020 northern hemisphere influenza vaccine^29^, exhibited only minimal cross reactivity against circulating viruses from the same clades, indicating that the circulating viruses differ antigenically from the vaccine strains of the same clade.

**Table 1.** Hemagglutination inhibition assay.

|  |  |  | Post-infection ferret sera raised against |  |  |  |  |  |  |
| --- | --- | --- | --- | --- | --- | --- | --- | --- | --- |
| Virus | Passage | Clade | NIB 104 | A/NL/314/2019 | NIB-112 | A/NL/3466/17 | A/NL/1802/18 | X-327 | A/NL/384/2019 |
|  |  |  | 3C.2a1a | 3C.2a1b | 3C.2a2 | 3C.2a2 | 3C.2a2 | 3C.3a | 3C.3a |
| A/Singapore/INFH-16-0019/16 (NIB 104) | E7Mdck_Siat2hCK | 3C.2a1a | 640 | 320 | 960 | 160 | 480 | <5 | 60 |
| A/Netherlands/1797/17 | Mdck1Siat2hCK | 3C.2a1 | 15 | 160 | 30 | 40 | 80 | <5 | 30 |
| A/Netherlands/314/2019 | MdckSiat2hCK | 3C.2a1b | 30 | 240 | 30 | 60 | 80 | <5 | 40 |
| A/Netherlands/10009/2019 | mixhCK2 | 3C.2a1b | 20 | 160 | 40 | 120 | 120 | <5 | 30 |
| A/Netherlands/371/19 | MdckSiatmixhCK | 3C2.a1b | <5 | 160 | 5 | 30 | 40 | <5 | 15 |
| A/Switzerland/8060/17 (NIB-112) | E7E1hCK | 3C.2a2 | 640 | 120 | 1920 | 1280 | 240 | <5 | <5 |
| A/Netherlands/3466/17 | Siat1hCK1 10-2 | 3C.2a2 | 15 | 80 | 320 | 480 | 80 | <5 | 30 |
| A/Netherlands/1802/18 | Siat1hCK1 10-1 | 3C.2a2 | 15 | 80 | 40 | 40 | 640 | <5 | 20 |
| A/Netherlands/10616/19 | MdckSiatmixhCK | 3C.2a2 | 15 | 80 | 320 | 640 | 80 | <5 | 20 |
| A/Kansas/14/17 (X-327) | E17E1Siat1hCK | 3C.3a | 640 | <5 | <5 | <5 | <5 | 320 | 60 |
| A/Netherlands/384/2019 | Siat2hCK 10-2 | 3C.3a | 30 | 30 | 60 | 40 | 80 | 40 | 240 |
| A/Netherlands/10006/2019 | MdckSiatmixhCK2 | 3C.3a | <5 | <5 | 20 | 20 | 30 | <5 | 240 |

The results of the HI assay were compared with a focus reduction assay (FRA) using the same sera and viruses (Table S2) ^30^. In this assay, the ability of antibodies to block virus infection in mammalian cell culture (MDCK-Siat cells, which overexpress Sia(α2,6)Gal moieties) is quantified. The FRA confirmed the trends observed in the HI assay (Fig. S7), indicating that the modified erythrocytes are reliable for antigenic characterization of A/H3N2 viruses.

### Molecular dynamics simulations of HAs in complex with their receptors

It is widely accepted that *α*2,6-sialyl LacNAc is the prototypic human receptor for IAVs^11^. All viruses that we examined, including A/H1N1 viruses (Fig. S8), recognized with high avidity *N*-glycans having an *α*2,6-sialoside on an extended LacNAc moiety. Furthermore, human respiratory tissue abundantly expresses such extended structures ^20^. Thus, we reasoned that mutational changes in recent A/H3N2 viruses led to a reduced binding affinity of the prototypic human receptor, which has been compensated by making interactions with an extended LacNAc chain, which are abundantly present in human respiratory tissue.

X-ray crystal structures ^1,31,32^ have shown that sialic acid is recognized in a conserved hydrophobic pocket (Y98, H183, Y195 and W153) (Table S3). It makes further interactions through a hydrogen bonding network with residues 135-137, E190 and S228. Sequence alignments showed that post-2000 strains acquired single point mutations at one of these residues (Table S3), which disrupted the hydrogen bonding network, and likely resulted in a reduced binding affinity of the prototypic human receptor^1^. Furthermore, HAs of early human A/H3N2 strains have Glu at residue 190, which can form a hydrogen bond with O9 of sialic acid, whereas post-2000 strains, favor Asp190 at this position, which due to its shorter side chain, cannot form such an interaction. The 190E/D mutation was accompanied by a 225G/D mutation, which resulted in a rotation of Gal-2 allowing a hydrogen bond interaction with the site chain of 225D ^1,31^. This rotation places an extended glycan chain closer to the 190-helix and potentially allows for addition interactions. To examine this mode of binding, we compared molecular dynamics generated structures of *α*2,6-sialyl poly-LacNAc in complex with NL91, NL03, and NL17 (Fig. 3). It recapitulated observations made by X-ray crystallography studies and provided insight in how mutational changes allowed for interactions with an extended LacNAc moiety. The MD trajectory of NL91 only showed interactions with sialic acid, including the important hydrogen bond between Sia-1 O9 and Glu190 (Fig. 3A)^1,31^. In the case of NL03 and NL17, Asp190 is at a distance of 4.5 Å to Sia-1 O9, and thus cannot establish a hydrogen bond (Fig. 3D and G). The G225D substitution resulted in a rotation of the bond between the sialic acid and Gal-2 to form an H-bond with its O3 ^10^, (Fig. 3E and 3H). As anticipated, the resulting rotation of Gal-2 placed the extended LacNAc moieties near the 190-helix ^32^ resulting in a hydrogen bond between Asp190 and Gal-4 O2, while either Ser or Asn193 provided a H-bond with the acetamide moiety of GlcNAc-3 (Fig. 3E). In addition, Gal-6 (Fig. 3E and 3H) makes a CH-*π* interaction with the side chain of Y159 (Fig. S10). Such interactions are often observed in glycan-protein complexes and contribute substantially to binding^33^. Structural analysis of the HAs of NL91 and NL03 showed that mutations distal to the receptor binding domain (A131T, H155T and E156H), reorient the side chain of Y159 resulting in an extended receptor binding site allowing interactions with Gal-6 (Fig. S9). The MD simulations support that A/H3N2 of the 3C.2 clade have undergone mutations to create and extended binding site to compensate for reduced binding of the prototypic human receptor (Fig. 3C, F and I). Interestingly, 3C.3a viruses, which can utilize shorter receptors having two LacNAc units (Fig. 1A), have a tyrosine to serine mutation at position 159 (Table S3). Thus, the dependence on extended receptors is reversible, and these viruses have found a way to bind shorter structures with sufficient affinity for infection. The observation that distant mutations can alter the position of a side chain of an amino acid in the receptor binding site, which subsequently can become susceptible to antigenic pressure, indicates that such remote mutations need to be considered for the evolution of receptor binding and antigenic distance.

**Figure 3.**
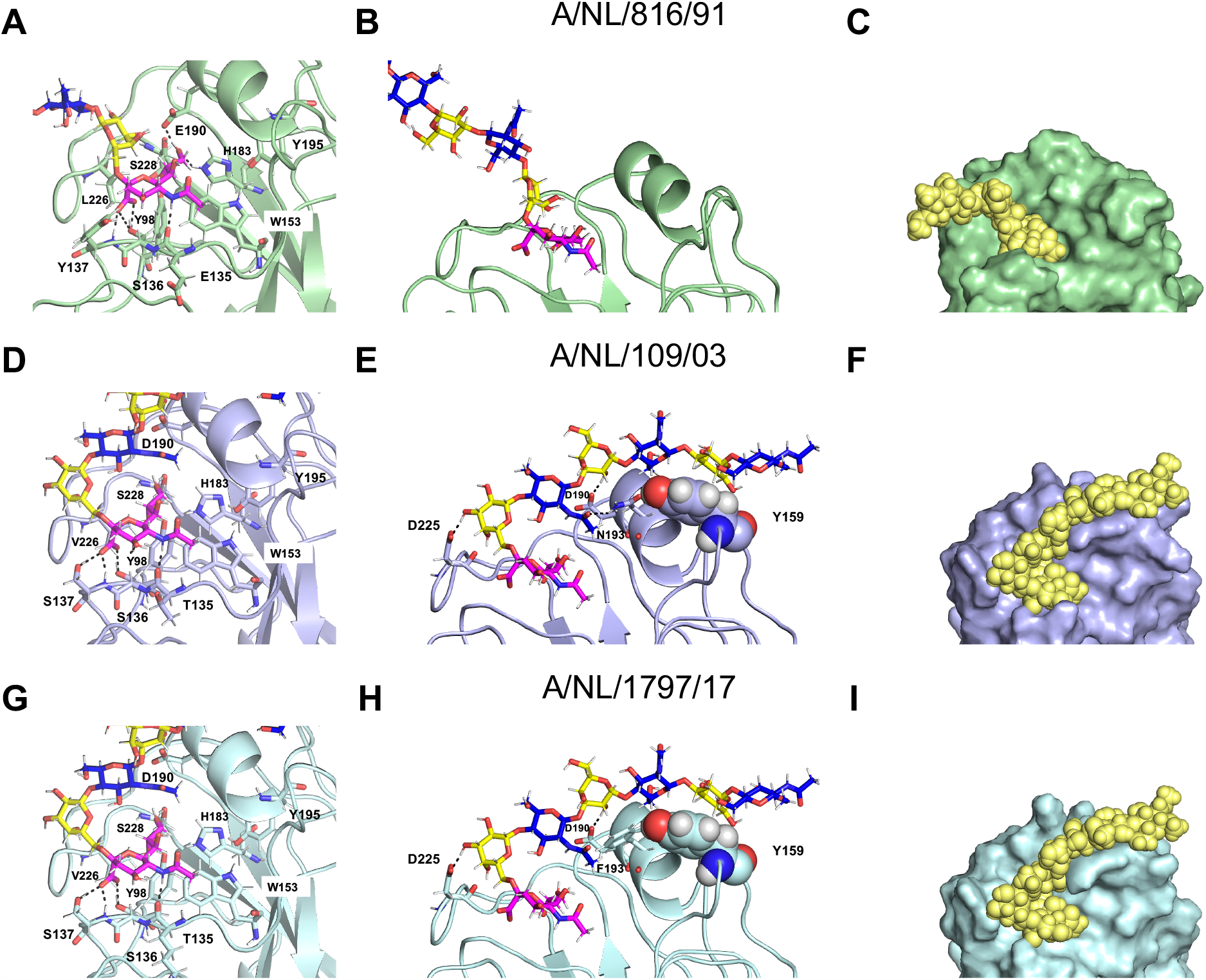
Structural comparison of HAs from evolutionary different strains in complex with extended glycan receptor. Details of the sialic acid binding sites are shown for A/NL/816/91 (**A**), A/NL/109/03 (**D**) and A/NL/1797/17 (**G**). The interactions of the poly-LN chain with the protein are shown for A/NL/816/91 (**B**), A/NL/109/03 (**E**) and A/NL/1797/17 (**H**). The surface and spheres representations of HA/glycans complexes are shown for A/NL/816/91 (**C**), A/NL/109/03 (**F**), and A/NL/1797/17 (**I**).

## Discussion

Understanding at a molecular level the evolution of receptor specificities and associated antigenic changes will offer important opportunities to address challenges in surveillance and vaccine production for A/H3N2 viruses. Here, such knowledge was exploited to engineer the surface of erythrocytes with functional receptors allowing easy antigenic characterization of recent A/H3N2 isolates using the HI assay. It confirmed that antigenically distinct viruses are circulating in humans and that egg-passaged A/H3N2 vaccine components match poorly to circulating strains. Due to the failure of the classical HI, focus reduction assays have been employed to antigenically characterize A/H3N2. These assays are, however, time-consuming, have low throughput, suffer from low reproducibility, and can lead to adapted mutations thereby providing incorrect results ^34^. These limitations are addressed by the glyco-engineered red blood cells described here. The altered receptor requirement of A/H3N2 viruses is also complicating propagation in the laboratory. Recently, an MDCK cell line (hCK) was introduced in which *α*2,3-sialyl transferases were genetically removed and ST6Gal1, which introduces human type receptors, was overexpressed ^24^. This cell line supports efficient replication of contemporary human A/H3N2 viruses that maintain higher genetic stability ^35^. The minimal receptor requirements of contemporary A/H3N2 viruses established in this study support further engineering of such cells to biosynthesize extended *α*2,6-sialylated LacNAc moieties. Furthermore, it is the expectation that the described glycan engineering approach for erythrocytes can be extended to other cells, such as MDCK cells, to quickly provide laboratory hosts for virus replication. Finally, an understanding of the evolution of receptor specificities of A/H3N2 viruses and associated antigenic changes will facilitate the development of predictive evolutionary models for the reliable selection of vaccine strains.

## Methods

### Virus production

#### Materials

Eagle’s minimal essential medium (EMEM), penicillin, streptomycin, L-glutamine, sodium bicarbonate, HEPES, 1x non-essential amino acids and N-tosyl-L-phenylalanine chloromethyl ketone (TPCK) treated trypsin were purchased at Lonza Benelux BV, Breda, The Netherlands. Fetal bovine serum was obtained from Greiner.

Madin-Darby canine kidney (MDCK), MDCK-Siat and hCK cells were cultured in Eagle’s minimal essential medium supplemented with 10% fetal bovine serum (FBS), 100 U mL^-1^ penicillin (P), 100 U mL^-1^ streptomycin (S), 2 mM L-glutamine (L-glu), 1.5 mg mL^-1^ sodium bicarbonate (NaHCO_3_), 10 mM HEPES, and 1x non-essential amino acids (NEAA) ^24^. In addition, hCK cells were supplemented with 2 µg mL^-1^ puromycin and 10 µg mL^-1^ blasticidin and MDCK-Siat cells were supplemented with 1 mg mL^-1^ Geneticine. To produce virus stocks, cells were washed twice with PBS 1 h after inoculation with the virus of interest and cultured in infection media, consisting of EMEM supplemented with 100 U mL^-1^ penicillin, 100 μg mL^-1^ streptomycin, 2 mM glutamine, 1.5 mg mL^-1^ sodium bicarbonate, 10 mM Hepes, 1x non-essential amino acids, and 20 μg mL^-1^ N-tosyl-L-phenylalanine chloromethyl ketone (TPCK) treated trypsin. To produce virus stocks in eggs, 100 μL of virus was inoculated in the allantoic cavities of 11-day-old embryonated hens’ eggs. The allantoic fluid was harvested after 2 days.

### Microarray studies

#### Materials

Virus isolates were produced as described above. Oseltamivir was purchased from Sigma Aldrich [Cat# SML1606]. CR8020 A/H3N2 stem antibody was kindly provided by Dr. Dirk Eggink and expressed following previously published procedures ^36^. Goat anti-human Alexa-647 [Cat# A21445] and streptavidin-AlexaFluor 635 [Cat# SA1011] antibodies were obtained from Thermo Fisher. Control lectins *Erythrina cristagalli* agglutinin (ECA) [Cat#B-1145], *Sambuca nigra* agglutinin (SNA) [Cat# B-1305], *Maackia amurensis* lectin I (Mal-I) [Cat# B-1315] were purchased from Vector Labs.

#### Arrayer and printing surfaces

Compounds were printed on amine reactive, NHS activated glass slides (NEXTERION ® Slide H) from Schott Inc using a Scienion sciFLEXARRAYER S3 non-contact microarray printer equipped with a Scienion PDC80 nozzle (Scienion Inc). Glycans were dissolved in printing buffer (sodium phosphate, 250 mM, pH 8.5) at a concentration of 100 µM. Each compound was printed in replicates of 6 with a spot volume of ∼400 pL, at 20 °C and 50% humidity. Slides were blocked with 5 mM ethanolamine in Tris buffer (pH 9, 50 mM) for 1 h at 50 °C and rinsed with DI water after printing.

#### Glycan microarray

Quality control was performed using the plant lectins ECA (specific for terminal Gal), SNA (specific for α2,6-linked Neu5Ac) and MAL-I (specific for α2,3-linked Neu5Ac) and is shown in Fig. S1. Quality control of the CR8020 A/H3N2 influenza hemagglutinin stem specific antibody specificity was performed by incubation of the antibody to the array as described below, in the absence of a virus (Fig. S1). The printed library of compounds comprised the glycans described in the Supplementary Information (#8, #9, #11, #12, #13, #15, #16, #17) and published previously (#1-#7, #10, #14) ^37^.

#### Sample

Virus isolates (25 µL) were diluted with PBS-T (PBS + 0.1% Tween, 25 µL) and applied to the array surface in the presence of oseltamivir (200 nM) in a humidified chamber for 1 h, followed by successive rinsing with PBS-T (PBS + 0.1% Tween), PBS and deionized water (2x) and dried by centrifugation. The virus-bound slide was incubated for 1 h with the CR8020 A/H3N2 influenza hemaglutinin stem specific antibody (100 µL, 5 µg mL^-1^ in PBS-T) and washed according to previous washing procedure. A secondary goat anti-human AlexaFluor-647 antibody (100 µL, 2 µg mL^-1^ in PBS-T) was applied, incubated for 1 h in a humidified chamber and washed again as described above. The control lectins containing a biotin tag were visualized with Streptavidin-AlexaFluor635. Slides were dried by centrifugation after the washing step and scanned immediately.

#### Detection and data processing

The slides were scanned using an Innopsys Innoscan 710 microarray scanner at the appropriate excitation wavelength. To ensure that all signals were in the linear range of the scanner’s detector and to avoid any saturation of the signals various gains and PMT values were employed. Images were analyzed with Mapix software (version 8.1.0 Innopsys) and processed with our home written Excel macro. The average fluorescence intensity and SD were measured for each compound after exclusion of the highest and lowest intensities from the spot replicates (n=4).

### Glycomic analysis

#### Materials

Acetic acid [Cat# 5330010050], MS grade formic acid [Cat# 5330020050], 2-aminoanthrallic acid (2-AA) [Cat#10680], dimethyl sulfoxide (DMSO) [Cat# W387520] and sodium cyanoborohydride [Cat# 156159] were obtained from Sigma Aldrich. Trifluoro acetic acid (TFA) was purchased from Acros Organics [Cat#434161000]. MS-grade acetonitrile (MeCN) was obtained from Biosolve [Cat# 200-835-2]. PNGase F was purchased from Roche Diagnostics (1U defined as the amount of enzyme catalysing the conversion of 1 µmol(substrate) min^-1^) [Cat#06538355103]. Denaturation buffer (0.5% SDS, 40 mM dithiothreitol (DTT)), 1% NP-40, and glycobuffer (50 mM sodium phosphate) were obtained from New England BioLabs. Ammonium formate was purchased from Fluka chemicals [Cat# AGG1946-85021], C18 solid phase extraction (SPE) Sep-Pak® Vac (1cc) columns from Waters Corporation [Cat# WAT054955], PGC SPE Hypercarb Hypersep (1cc) columns from Thermo Scientific [Cat# 60106-303] and PD Minitrap Sephadex G-10 size exclusion cartridges from GE Healthcare [Cat# 28-9180-10]. MilliQ water was obtained from a Synergy® water purification system.

#### N-glycan extraction and release from erythrocytes

Cell surface N-glycans were extracted according to a reported protocol ^26^. Briefly, erythrocytes (400 µL, 50%) were concentrated by centrifugation (430 rcf, 10 min) and removal of the supernatant. Erythrocytes were lysed under gentle shaking at room temperature for 1 h using deionized water (3x pellet size). The suspension was centrifuged (4000 rcf, 10 min), the supernatant removed, and the pellet resuspended in deionized water. The process was repeated until the pellet decolorized, indicating an efficient lysis of the erythrocytes. Denaturing of the cell membrane pellet was performed by heating for 10 min to 95 °C in denaturation buffer (0.5% SDS, 40mM DTT in H_2_O). The N-glycans were released during overnight incubation at 37 °C with PNGase F (5 U) in a sodium phosphate buffer (50 mM, pH 7.5) or with Endo F2 in sodium acetate buffer (50 mM, pH 4), both containing NP-40 (1%).

#### Purification and labeling of N-glycans

The released N-glycans were applied to a C18 SPE cartridge and glycans were eluted with 5% MeCN in H_2_O (0.05% TFA, 1 mL). The eluate was further purified on a PGC SPE cartridge by gradually increasing the hydrophobicity from 100% H_2_O (0.05% TFA, 1 mL) to 5% MeCN in H_2_O (0.05% TFA, 1 mL), and to 50% MeCN in H_2_O (0.05% TFA, 1 mL) which eluted the glycans. After drying in a N_2_ flow, the sample was dissolved in H_2_O (10 μL) and labelled by addition of a solution (10 μL) of 2-AA (48 mg mL^-1^) and sodium cyanoborohydride (63 mg mL^-1^) in DMSO/acetic acid (10:3 v/v) for 2 h at 65 °C. The crude mixture was diluted with H_2_O (80 µL) and purified on a Minitrap Sephadex G-10 gravity column by washing with of H_2_O (700 µL) and eluting with H_2_O (600 µL). The eluate was dried under N_2_ flow and dissolved in 20 µL of 50% MeCN in H_2_O prior to LC-MS analysis.

#### HILIC-IMS-QTOF analysis of N-glycans and data treatment

The *N*-glycan analysis was performed on a 1260 Infinity liquid chromatography system (Agilent Technologies) coupled to a 6560 IM-QTOF mass spectrometer (Agilent Technologies) and HPLC separation with a ZIC®-cHILIC column (3 µm, 100 Å, 150 mm × 2.1 mm, Merck) and a similar guard column (20 mm × 2.1 mm, Merck) using a gradient from 30% A to 50% A within 25 min (A: 50 mM (NH_4_)_2_CO_3_ in H_2_O; B: MeCN; 0.2 mL min^-1^; 60 °C). MS analysis was performed with a drying gas temperature of 300 °C and a flow of 8 mL min^-1^. The nebulizer pressure was set to 40 psi, the sheath gas flow to 11 L min^-1^ and the temperature to 350 °C. Measurements were run in negative mode with the capillary established at 3500 V. During the runs the Agilent tuning mix was infused for mass calibration based on reference signals at m/z 112.9855 and m/z 1033.9881.

The data analysis was performed with the Mass Hunter IM-MS Browser and the find feature function filtering for masses with an ion intensity of ≥500. Found masses were processed with the online Glycomod tool to identify glycan related masses ^38^.

### Glyco-engineering of erythrocytes

#### Materials

*Arthrobacter ureafaciens* neuraminidase was purchased at New England Biolabs (1 U defined as the amount of enzyme catalyzing the conversion of 1 µmol(substrate) min^-1^) [Cat# P0722L]. Mammalian glycosyltransferases were expressed according to literature reports ^28^. B3GnT2 and ST6Gal1 were cleaved and purified from GFP tags prior to use. Alkaline phosphatase (FastAP) was purchased at Thermo Scientific [Cat# EF0651]. Nucleotide sugars UDP-Gal, UDP-GlcNAc and CMP-Neu5Ac were obtained from Roche Diagnostics [UDP-Gal: Cat# 07703562103; UDP-GlcNAc: Cat# 06369855103; CMP-NeuAc: Cat# 05974003103].

#### Erythrocyte preparation

Fresh blood from chicken or turkey was centrifuged (10 min, 430 rcf) followed by removal of the supernatant. Pellets were washed three times in PBS with intermittent centrifugation (430 rcf, 10 min). Erythrocyte solutions were stored in a 50% solution in PBS until further use.

#### Enzymatic extension

To a suspension of fowl erythrocytes (250 µL, 50%), PBS (900 µL) and *Arthrobacter ureafaciens* neuraminidase (12 U) were added. The cells were incubated for 6 h at 37°C while tilting. Next, glycosyltransferases B4GALT1 (37,5 µL, 1 mg mL^-1^) and B3GnT2 (37,5 µL, 1 mg mL^-1^), the nucleotide sugars UDP-Gal (4.4 mM) and UDP-GlcNAc (4.4 mM), alkaline phosphatase (6 U), MnCl_2_ (2 mM) and BSA (6 µL, 2 mg mL^-1^) were added. This reaction mixture was incubated overnight at 37 °C while tilting. The erythrocytes were washed in PBS (2x, 600 µL) and the pellet was reconstituted in 900 µL PBS. Resialylation of the erythrocytes was performed using ST6Gal1 (37,5 µL, 1 mg mL^-1^) and CMP-Neu5Ac (4.4 mM) in the presence of alkaline phosphatase (6 U) and BSA (6 µL, 2 mg mL^-1^) for 4 h at 37 °C while tilting. The erythrocytes were washed in PBS (1x, 600 µL) and diluted to a 1% solution for hemagglutination (inhibition) assays.

### Stability assay

Fresh fowl erythrocytes were glyco-engineered as described above or used unmodified. A hemagglutination assay was performed every two days using two viruses, A/NL/761/09 and A/NL/816/91. Hemagglutination assays were performed as described below in full biological triplicates. The means were plotted ± SEM and are shown in Fig. S6.

### Hemagglutination assay

Hemagglutination assays were performed following standard procedures ^8^. Briefly, virus stocks were two-fold serial diluted in the presence of oseltamivir (20 nM). Turkey erythrocytes (1%, 25 µL) were mixed with the serial diluted viruses and incubated for 1 h at 4 °C before recording of the results. Titers were expressed as the highest dilution of virus stock that completely agglutinated the turkey erythrocytes.

### Hemagglutination inhibition assay

Hemagglutination inhibition assays were performed following standard protocols ^8^. Briefly, for the preparation of the antisera, ferrets were inoculated intranasally and blood was obtained 14 d later. Antisera were pre-treated with receptor destroying enzyme (RDE) by incubating overnight with an in-house produced filtrate of *Vibrio cholera* at 37 °C followed by 1 h incubation at 56 °C. The treated antisera were pre-absorbed with extended turkey erythrocytes (10%) in two cycles of 1 h incubation at 4 °C. Pre-absorbed antisera were two-fold serial diluted (starting at 1:10) and mixed with virus stock (25 µL) containing 4 hemagglutinating units. Viruses were incubated with the antisera for 1 h at 4 °C in the presence of BSA (0.25%) and oseltamivir (20 nM). Turkey erythrocyte solution (25 µL, 1%) was added and after 1 h incubation at 4 °C inhibition patterns were recorded. Titers were expressed as the value of the highest serum dilution that gave complete inhibition of agglutination.

### Focus reduction assay

Focus reduction assays were performed following standard protocols ^39^. First, infectious titers of the virus stocks were determined in hCK cells as described previously ^40^. RDE-treated sera were two-fold diluted in a 96-well plate (starting at 1:10) and mixed 1:1 with 100 TCID_50_/50 µL of virus. After 1 h incubation at 35 °C, 100 µL of the mixtures were transferred to hCK cells and after 90 min incubation at 35 °C, cells were washed and overlaid with 1.6% carboxymethylcellulose. After 48 h at 35 °C, cells were washed and fixed with formalin and permeabilized using 0.5% Triton X-100 for 10 min at room temperature. Subsequently, immunostaining was performed using a mouse monoclonal antibody (HB65; EVL, Woerden, The Netherlands) directed against the viral nucleoprotein (NP), followed by a horseradish peroxidase-labeled goat anti mouse immunoglobulin preparation (GAM-HRPO, Invitrogen, Foster city, CA), both for 1 h at room temperature. After washing, True-Blue substrate (KPL, Gaitherburg, Maryland) was added followed by a 10 min incubation at room temperature. The plates were washed, dried, and submitted to automated image capture using a Series 6 ImmunoSpot Image Analyzer (CTL Immuno-Spot, Cleveland OH, USA) to quantitate the percentage well area covered by spots of infected cells. Inhibition ≥ 90% was considered positive for neutralization.

### Structural studies

#### Alignment

To investigate the relevance of specific mutations in defining receptor preferences during antigenic drift, primary amino acid sequences were compared of A/H3N2 HAs from 1968 to 2019. Protein sequences and structures were derived from the 3DPlu database (http://3dflu.cent.uw.edu.pl/index.html)^41^ and the GISAID webpage (https://www.gisaid.org). It identified specific mutations that may influence receptor binding preferences. We paid specific attention to mutations in the four structural elements that define the RBS, including the 130-loop, the 150-loop, the 190-helix and the 220-loop. The results from this analysis are summarized in Table S3.

#### MD simulations

A starting pose of the NL91 from 1991 was generated by superimposition of the model derived structure (id code ACU12494) [http://3dflu.cent.uw.edu.pl/index.html] onto the X-ray crystal structure of the A/HK/1/1968 H3N2 influenza virus hemagglutinin in complex with 6’-SLNLN (pdb code 6TZB) ^32^. The starting pose of NL03 was generated by using the X-ray crystal structure of the A/Wy/3/03 influenza virus hemagglutinin in complex with 6’-SLN (pdb code 6BKR)^31^. Glycan receptors, NeuAc*α*2-6(LacNAc)_2_ and NeuAc*α*2-6(LacNAc)_3_, were generated by using the carbohydrate builder GLYCAM-web site [http://glycam.org]. The glycosidic torsion angles of the monosaccharides were maintained as observed by X-ray crystallography, while those not resolved were defined according to the lower energy values predicted by the GLYCAM-web modeling tool. The structural model of the NL17 was obtained by using the mutagenesis tool implemented in PyMOL. The resulting poses were used as starting points for molecular dynamics (MD) simulations. The MD simulations were performed using the Amber16 program4 with the protein.ff14SB, the GLYCAM_06j-1 and the water.tip3p force fields parameters. Next, the starting 3D geometries were placed into a 10 Å octahedral box of explicit TIP3P waters, and counter ions were added to maintain electroneutrality. Two consecutive minimization steps were performed involving (1) only the water molecules and ions and (2) the whole system with a higher number of cycles, using the steepest descent algorithm. The system was subjected to two rapid molecular dynamic simulations (heating and equilibration). The equilibrated structures were the starting points for a final MD simulations at constant temperature (300 K) and pressure (1 atm). 100 ns Molecular dynamics simulations without constraints were recorded, using an NPT ensemble with periodic boundary conditions, a cut-off of 10 Å, and the particle mesh Ewald method. A total of 50 000 000 molecular dynamics steps were run with a time step of 1 fs per step. Coordinates and energy values were recorded every 50000 steps (50 ps) for a total of 1 000 MD models. The detailed analysis of the H-bond and CH-*π* interactions was performed along the MD trajectory using the cpptraj module included in Amber-Tools 16 package.

## Acknowledgments

We thank T. Manders, dr. H.G.R. Matthijs and dr. R.M. Dwars (Faculty of Veterinary Sciences, Utrecht University) for providing chicken blood and M. Pronk and R. van Beek (Department of Viroscience, Erasmus MC) for technical assistance. Dr. L. Liu (CCRC) and dr. M.A. Wolfert (Utrecht University) developed, printed and validated the glycan microarray. We would like to thank Dirk Eggink from the Amsterdam Medical Center for supplying the CR8020 antibody.

## Author contributions

G.-J.B., R.P.dV., F.B. and R.J.vB. designed the project; R.J.vB., F.B., L.U., T.B., and R.P.dV. performed experiments and data analysis; R.A.F., G.-J.B., S.H. and R.P.dV. provided scientific guidance on experimental setup and data interpretation; K.W.M., D.C., and J.Y.Y. provided recombinant enzymes; G.-J.B., R.J.vB., F.B., L.U., S.H., R.A.F. and R.P.dV. wrote the manuscript; and all authors provided comments and suggestions on the manuscript.

## Data and materials availability

All data is available in the main text or the supplementary materials. Sharing of materials described in this work will be subject to standard material transfer agreements.

